# Glycome profiling and immunohistochemistry uncover spaceflight-induced changes in non-cellulosic cell wall components in *Arabidopsis thaliana* seedling roots

**DOI:** 10.1101/2023.03.14.532448

**Authors:** Jin Nakashima, Sivakumar Pattathil, Utku Avci, Sabrina Chin, J. Alan Sparks, Michael G. Hahn, Simon Gilroy, Elison B. Blancaflor

## Abstract

A large and diverse library of glycan-directed monoclonal antibodies (mAbs) was used to determine if plant cell walls are modified by low-gravity conditions encountered during spaceflight. This method called glycome profiling (glycomics) revealed global differences in non-cellulosic cell wall epitopes in *Arabidopsis thaliana* root extracts recovered from RNA purification columns between seedlings grown on the International Space Station-based Vegetable Production System and paired ground (1-*g*) controls. Immunohistochemistry on 11-day-old seedling primary root sections showed that ten of twenty-two mAbs that exhibited spaceflight-induced increases in binding through glycomics, labeled space-grown roots more intensely than those from the ground. The ten mAbs recognized xyloglucan, xylan, and arabinogalactan epitopes. Notably, three xylem-enriched unsubstituted xylan backbone epitopes were more intensely labeled in space-grown roots than in ground-grown roots, suggesting that the spaceflight environment accelerated root secondary cell wall formation. This study highlights the feasibility of glycomics for high-throughput evaluation of cell wall glycans using only root high alkaline extracts from RNA purification columns, and subsequent validation of these results by immunohistochemistry. This approach will benefit plant space biological studies because it extends the analyses possible from the limited amounts of samples returned from spaceflight and help uncover microgravity-induced tissue-specific changes in plant cell walls.

## INTRODUCTION

Land plants evolved under the continuous presence of the Earth’s gravitational field. As such, plant development on Earth is strongly influenced by gravity ^1^. One of the most obvious manifestations of gravity’s effects on plants is the redirection of the growth of their major organs when they are turned on their sides. Whereas roots redirect their growth downward toward the gravity vector, most shoots grow upward, opposite the pull of gravity. This process called gravitropism is widely regarded as a major environmental signal that shapes the architecture of roots and shoots, enabling plants to more efficiently capture soil resources and light inputs required for optimal growth ^2^.

In shoots, the initial growth redirection of the organ due to gravitropism is followed by a sustained and upward growth. The continuous growth of shoots opposite the direction of gravity requires mechanical strength provided by the rigid, polysaccharide-rich cell wall, which is a process referred to as gravity resistance and is independent of gravitropism ^3^. Signals for the gravity resistance response are believed to be perceived by mechanoreceptors that transmit information directing cortical microtubules to reorient. These changes in cortical microtubule orientation then lead to modified cell wall properties in the plant ^4^. Because of their importance in providing mechanical support for processes, such as gravity resistance, cell walls and the elaboration of the molecular machinery required for their construction and remodeling, played a major role in the transition of plants from an aquatic environment to life on land ^5^.

As the US National Aeronautics and Space Administration (NASA) considers the return of humans to the Moon, and for astronauts to set foot on Mars, the development of technological platforms for safe and sustainable space transportation will be essential. A crucial component for the creation of such technology is a convenient and reliable source of nutrient-rich food and oxygen for humans as they embark on long duration spaceflight missions ^6^. Plants fulfill this need as they not only provide food and oxygen, but are also known to help in recycling waste, removing CO2, and improving the psychological well-being of astronauts who have to spend months, and likely years living in a confined spacecraft environment ^7,8^. However, before plants can be deployed as major components of advanced life support systems, understanding how their biology is affected by the space environment is essential ^9–11^. Doing so will enable the development of plant cultivars that are better adapted to the unique environment of space, including reduced gravity (*i.e*., microgravity) and confined growth environments, both of which can influence gravitropism and gravity resistance.

The era of the Space Shuttle and the International Space Station (ISS) enabled a number of experiments that address how spaceflight influences plant growth from whole plant and cell/tissue responses, to changes in the expression of genes and proteins ^12–18^. In transcriptomic studies of spaceflight grown *A. thaliana* seedlings, genes that are often found to be differentially regulated by spaceflight are those encoding proteins related to cell wall remodeling and oxidative stress ^11,14,16,19^. Changes in the expression of cell wall-related genes may be due to the reduced mechanical load encountered by plants in microgravity. In fact, studies that directly measure cell wall metabolites or microscopically examine cell walls from plants grown in space are consistent with spaceflight transcriptomics data. For example, cellulose microfibrils of soybean seedlings grown on the Foton-M2 capsule are more disorganized than those of seedlings grown on Earth. Disorganized cellulose microfibrils of space-grown seedlings leads to thinner xylem vessel walls ^20^. Another study showed that expression of microtubule regulatory proteins and the percentage of cells with transverse cortical microtubules increases in *A. thaliana* seedlings grown in microgravity ^21^. Cortical microtubules serve as tracks for cellulose synthesizing complexes ^22^. Thus, reduced growth anisotropy of microgravity-grown seedlings, which had longer and thinner hypocotyls than those of Earth-grown seedlings, could be partly explained by cortical microtubule-influenced modifications in cellulose microfibril orientation ^21^.

Plant cell walls consist primarily of cellulose, hemicellulose, and pectin, and in secondary walls, lignin ^23,24^. The polysaccharide composition of each cell wall type varies among plant tissue and cell types, stage of plant development, and the environment, in which the plant is grown ^25–27^. Glycome profiling is a powerful tool to study the complexity of plant cell walls. It involves the use of a large and diverse collection of monoclonal antibodies (mAbs) that recognize unique epitopes in most major non-cellulosic plant cell wall glycans ^28^. The binding of mAbs to specific cell wall glycan epitopes as determined by Enzyme-Linked Immunoabsorbent Assay (ELISA), provides a comprehensive picture about environmentally- or genetically-induced changes in cell wall composition and structure ^28–30^. However, the power of glycome profiling is best realized when it is combined with immunohistochemistry, which enables the visualization of cell wall changes occurring in specific tissue and cell types ^29^. Here, as part of the <u>A</u>dvanced <u>P</u>lant <u>EX</u>periments (APEX) 03-1 in space project, we demonstrate the feasibility of combining glycomics and immunohistochemistry to study changes in non-cellulosic cell wall composition of *A. thaliana* roots adapting to spaceflight. We show that plant biomass recovered from RNA extraction columns provides valuable information on spaceflight-induced global changes in cell walls, some of which are manifested as tissue/cell-type differences in cell wall epitope accumulation between ground- and space-grown seedling roots.

## RESULTS

### Seedling morphology of space- and Earth-grown seedlings in the Vegetable Production System

Six and eleven days after experiment activation (Supplementary Fig. 1), seedlings from ground controls and space displayed robust growth on the ISS-based and ground Vegetable Production System (Veggie). Six-day-old seedlings had about two true leaves, in addition to fully expanded cotyledons (Fig. 1a, c). On the other hand, 11-day-old seedlings had 4 - 5 true leaves and root systems with extensive laterals (Fig. 1b, d). As reported previously, roots from seedlings grown in space skewed to one side, while those on the ground grew downwards in alignment with the gravity vector ^12,18,31^. The directional skewing of primary roots was also observed in 6-day-old APEX 03-1 seedlings grown in space (Fig. 1c). In 11-day-old space-grown seedlings, many roots grew toward the shoot with some roots reaching the top of the plate (Fig. 1d). Unlike roots, there were no clear qualitative differences in growth characteristics between shoots of ground- and space-grown seedlings. Therefore, we focused on characterizing root responses. To maximize return from the limited samples from spaceflight, we took theRNA*later*-fixed root debris left over from RNA extraction columns used in a separate transcriptomic study of these plants for glycome profiling. A separate set of seedling roots fixed in aldehydes were used for immunohistochemistry.

**Fig. 1.**
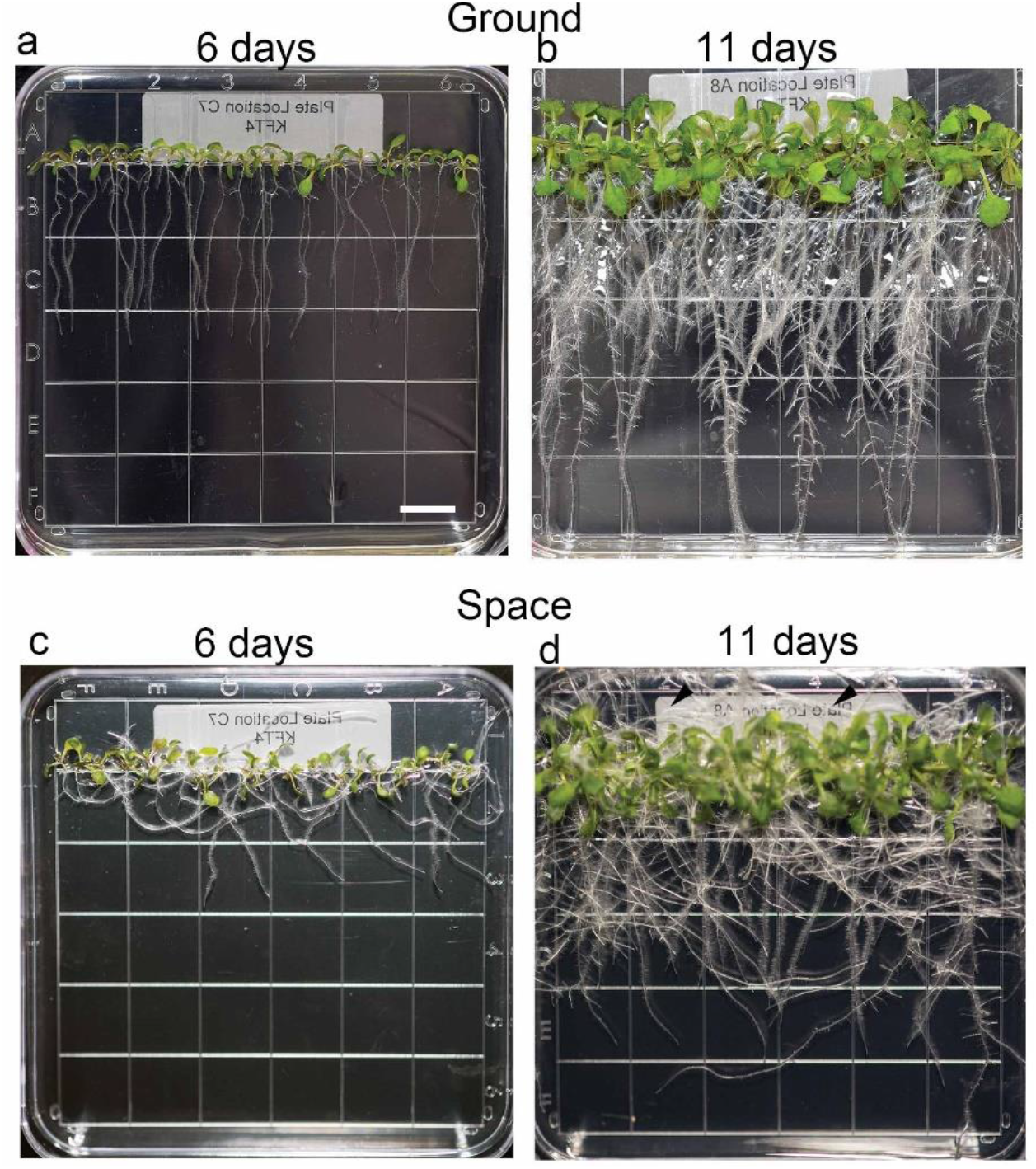
Roots of space and Earth-grown *Arabidopsis thaliana* seedlings on square Petri dishes in Veggie differ in growth direction. Representative images of six- (a and c) and 11-day-old (b and d) seedlings prior to harvest and fixation. Note that roots of seedlings on the ground grow vertically toward the direction of gravity (a, b). On the other hand, roots of 6-d-old seedlings in space skewed generally to the right (c). Some roots of 11-d-old seedlings reached the top of the plate (arrowheads). Bar = 1 cm.

### Glycome profiling reveals global differences in glycan-directed monoclonal antibody binding to root cell wall extracts between space- and Earth-grown seedlings

Glycome profiling involves sequential extraction of cell wall samples, consisting of six steps with increasingly harsh reagents in the following order: ammonium oxalate, sodium carbonate, 1 M potassium hydroxide (KOH), 4 M KOH, acidic sodium chlorite, and 4 M KOH post chlorite ^32^. A complete glycome profile analysis with six extraction steps normally requires 50-200 mg of cell wall material, which is roughly equivalent to 0.5-2 grams of plant biomass ^32^. For the glycomics experiments described here, the amount of plant material was a limiting factor. The RNA*later*-fixed seedlings were used for RNA-Seq studies, but we realized that the plant debris retained in RNA extraction columns was from the cell walls of the samples, and therefore could potentially be used for glycomics to assess spaceflight-induced global changes in cell walls. Seedling root and shoot debris from QIAGEN RNeasy Mini Kit RNA extraction columns yielded approximately 2 mg of cell wall material. The low amount of material allowed the use of only a single extraction step (4 M NaOH), rather than the six sequential fractionations noted above. The fourth 4 M KOH extraction step was chosen because it can yield the highest amount and diversity of alkaline soluble cell wall glycans, and therefore should provide the most information on cell wall composition differences between ground- and space-grown seedlings.

Glycome profiling with the 4 M KOH extract revealed that xylans and xyloglucans, which are major hemicelluloses in dicotyledonous plant cell walls ^23^, were the most abundant carbohydrates detected in root extracts of ground controls and spaceflight samples. This conclusion was based on the higher binding intensities of xylan- and xyloglucan-directed mAbs than those of mAbs directed to other non-cellulosic polysaccharide epitopes (Fig. 2a; Supplementary Table 1). Signal intensity of mAb binding was also generally higher in extracts of roots from space-grown seedlings than from extracts obtained from Earth-grown seedlings (Fig. 2b, c). Xylan mAbs with higher binding intensity included those that recognized small degrees of polymerization (DP) in the unsubstituted xylan backbone (*e.g*., DP5 to 8; CCRC-M138, CCRC-M139, CCRC-M140, and CCRC-M148) and those with methylglucuronic acid (MeGlcA-Xylan; CCRC-M144) and arabinose (Ara) side chains (CCRC-M108) (Fig. 2b). Xyloglucan epitopes included those that were xylosylated (Xyl-XG; CCRC-M100), galactosylated (Gal-XG-2; CCRC-M55, CCRC-M96, and CCRC-M50), and fucosylated (Fuc-XG; CCRC-M1 and CCRC-M84). Signal intensity of mAb binding was generally higher in root extracts from 11-day-old seedlings than that of root extracts from 6-day-old seedlings (Fig. 2a). Furthermore, mAb binding was more consistent among three biological replicates in root extracts from 11-day-old seedlings than root extracts from 6-day-old seedlings, which prompted us to focus our immunohistochemical studies on the older seedlings.

**Fig. 2.**
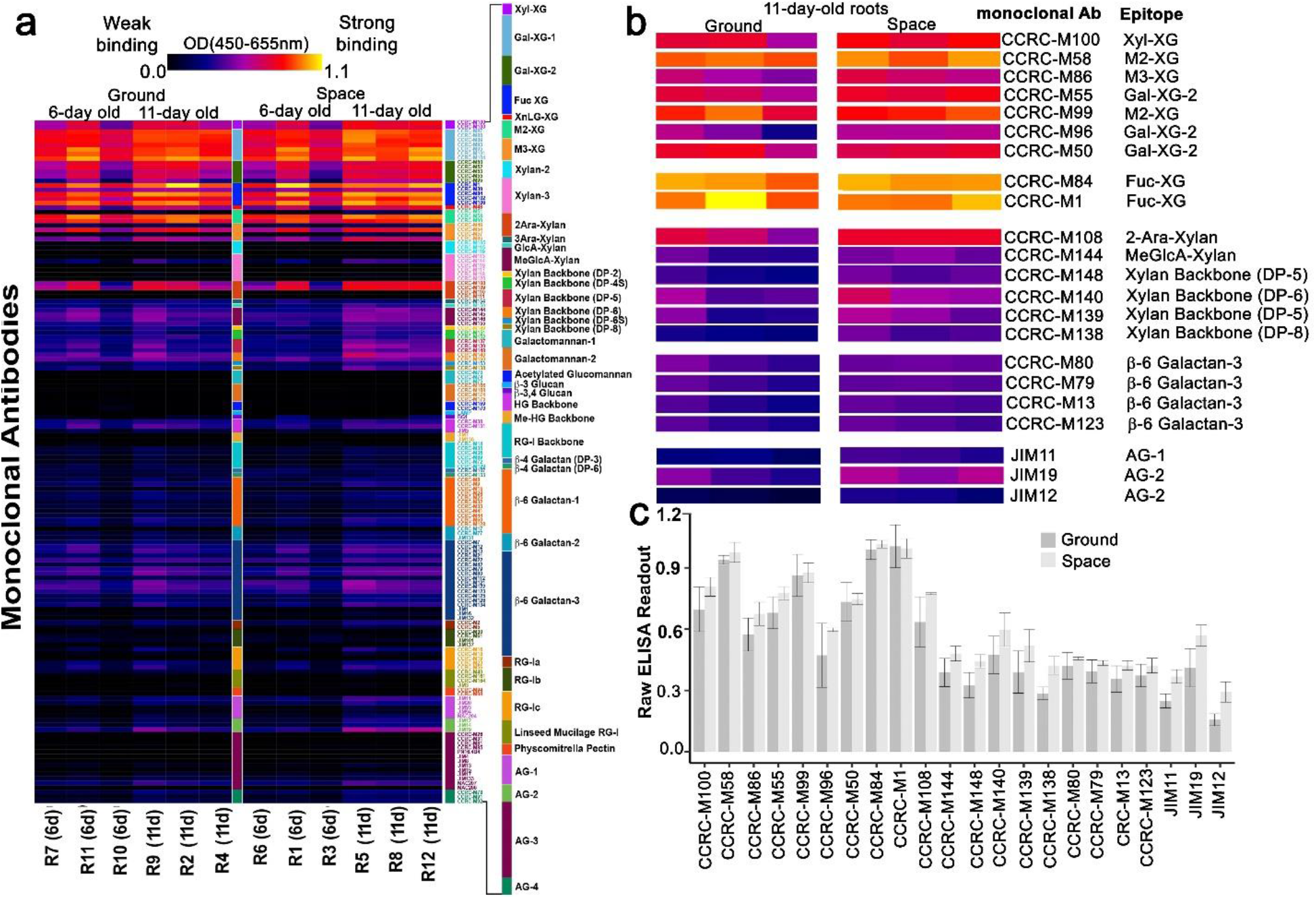
Glycome profiling reveals global differences in mAb binding to non-cellulosic cell wall epitopes between of root extracts of seedlings grown in space and those on Earth. (a) ELISA heat map of 4 M KOH root extracts from 6- and 11-day-old space-grown and ground control seedlings. (b) ELISA heat map of 11-day-old roots of 22 mAbs selected for immunohistochemistry. The mAbs selected bind to xylosylated xyloglucan (Xyl-XG), galactosylated xyloglucan (Gal-XG) fucosylated xyloglucans (Fuc-XG), and larger not yet defined xyloglucan epitopes, α-1,2-linked arabanosyl xylan (2-Ara-Xylan), 4-O-Me-GlcA xylan, unsubstituted xylan backbones (DP5-8), β-6-linked galactans, and arabinogalactan (AG) epitopes. (c) Bar graph of raw ELISA readout values of the 22 mAbs shown in panel (b). Values are means (n=3) ± Standard Deviation. Heat maps and bar graphs were generated from ELISA values in Supplementary Table 1. Galactose (Gal); Fucose (Fuc); Arabinose (Ara); Methyglucoronic acid (MeGlcA); DP (Degree of Polymerization); Xylose (Xyl).

Because glycome profiling of the debris from the RNA isolation columns revealed global differences in xyloglucan levels between roots from ground- and space-grown seedlings (Fig. 2), procedural controls using plant debris from xyloglucan xylosyltransferase (*xxt1/xxt2*) double and *mur3-3* single mutants were conducted. The *xxt1/xxt2* mutant has no xyloglucan ^33^ while the *mur3-3* mutant, which is defective in a gene encoding a galactosyltransferase, contains xyloglucans that lack a galactose-fucose side chain ^34,35^. Both mutants were stored in RNA*later* following the same timeline as the APEX 03-1 pipeline (Supplementary Fig. 1), and were processed in a similar manner as seedlings returned from space. Results showed that the binding of mAbs to a range of xyloglucan epitopes was significantly lower or absent in extracts of the *xxt1/xx2* mutants than that in extracts of wild-type seedlings. In *mur3-3* plant extracts, mAb binding to fucosylated xyloglucan was lower than that of wild type (Supplementary Fig. 2). These results are consistent with previously published data on these mutants that were obtained from cell wall samples prepared according to standard six-step extraction glycome profiling protocols rather than using debris from RNA purification columns, thus validating our results from the space-grown plants.

### Immunohistochemistry uncovers spaceflight-induced tissue/cell-type specific changes in non-cellulosic root cell wall components

Although ELISA readouts for glycome profiling revealed global differences in non-cellulosic cell wall components between root extracts of space- and Earth-grown seedlings, immunohistochemistry could provide more detailed information on where such changes are occuring. Twenty-two mAbs that exhibited higher binding to root cell wall extracts of space-grown seedlings than to extracts from ground controls were selected for immunohistochemistry based on the raw ELISA readout (Fig. 2b, c). These twenty-two mAbs recognized diverse epitopes of xylans, xyloglucans, and arabinogalactans (AGs) (Fig. 2b).

Two regions of 11-day-old roots were selected for immunohistochemistry (Fig. 3a). One region was located close to the root-hypocotyl junction, which represented mature root tissue (Fig. 3b). The other region was from the apical three millimeters of the primary root tip, which represented actively dividing and expanding cells of the meristem and elongation zone, respectively (Fig. 3c). Cross and longitudinal sections (0.25 μm) for mature root tissues and primary root tips, respectively, were obtained for immunohistochemistry.

**Fig. 3.**
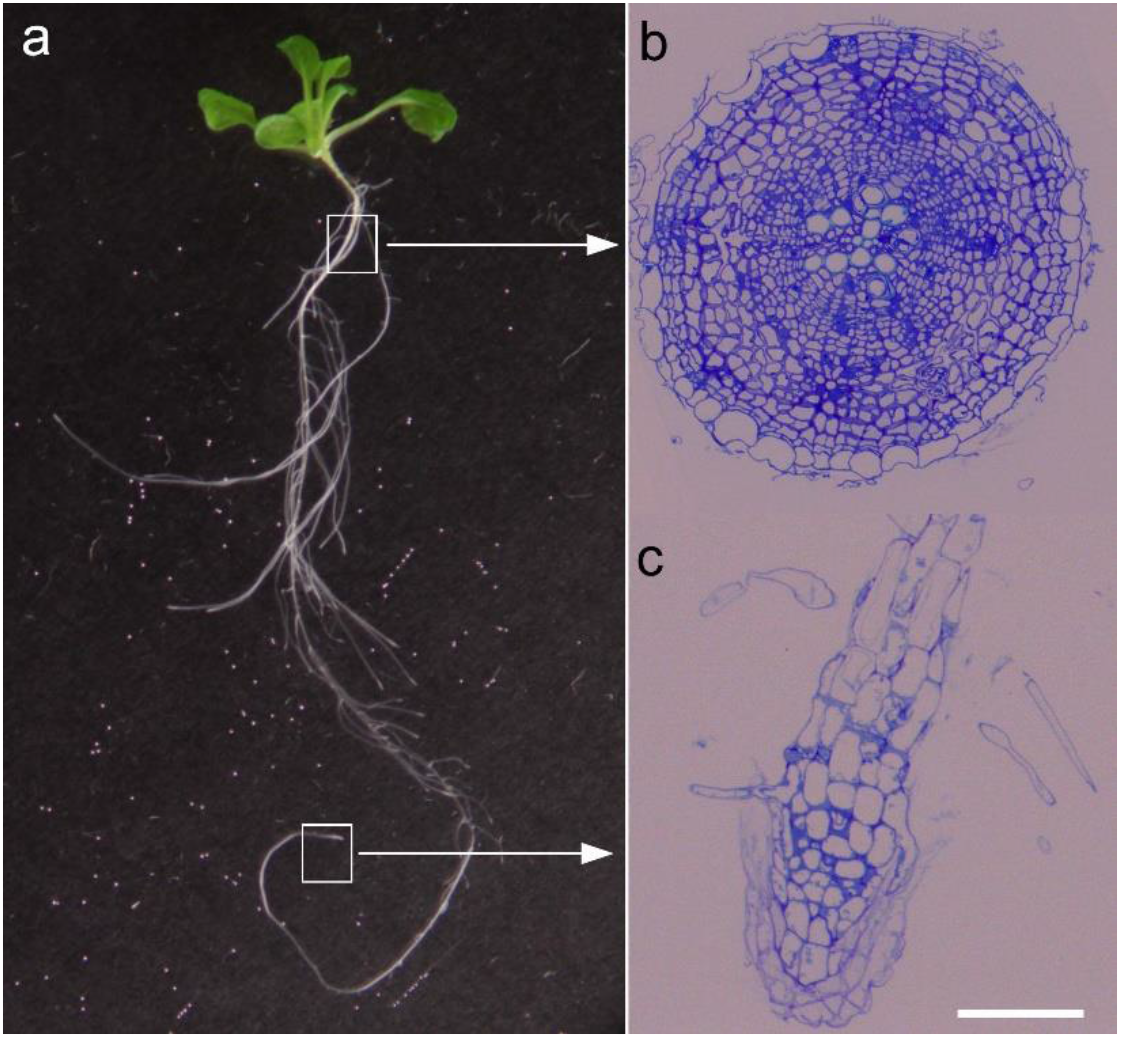
Regions of *A. thaliana* roots from 11-day-old seedlings used for immunohistochemistry. (a) A representative whole seedling grown in space. Representative toluidine Blue-O-stained semi-thin (0.25 μm thick) longitudinal section of the root tip (b) and cross-section of the root maturation zone (c) at the root-hypocotyl junction. Bar in (b) and (c) = 100 μm.

Immunohistochemistry of root tip longitudinal sections revealed that labeling intensity of 10 of the 22 mAbs was higher in space-grown seedlings than that in ground controls. Among the ten mAbs, eight mAbs recognized xyloglucan epitopes, one was specific to a xylan epitope with arabinose side chains (2-Ara-Xylan), and one was directed to a β-6 Galactan-3 epitope (Fig. 4a-f; Supplementary Fig. 3). Seven of the 22 mAbs that recognize xylan and AG epitopes did not label root tips from space- and Earth-grown seedlings (*i.e*., sections showed no fluorescence signals). Furthermore, CCRC-M80 and CCRC-M100, which recognize a β-6 Galactan-3 and Xyl-XG epitope, respectively, had fluorescence signals that were similar in intensity between roots of space- and Earth-grown seedlings (Supplementary Fig. 3). Two mAbs that also recognized β-6 Galactan-3 epitopes (CCRC-M79 and CCRC-M123) and an AG-2 epitope (JIM19) showed lower root fluorescence signals in space than those in the ground controls (Fig. 5a-f).

**Fig. 4.**
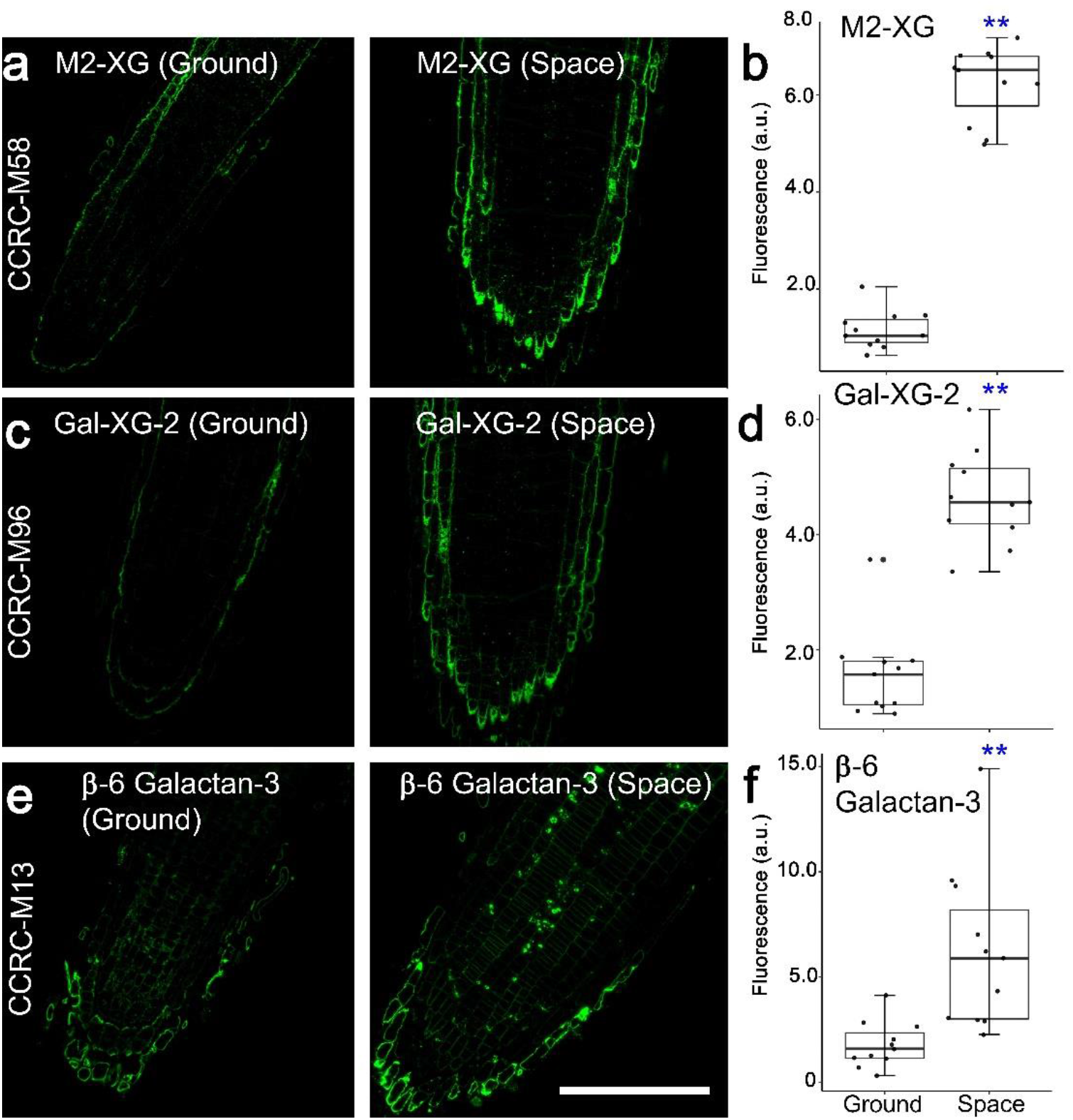
Immunohistochemistry of root tip longitudinal sections reveals spaceflight-induced increases in mAb binding to xyloglucan (XG) and β-6 Galactan-3 epitopes. M2-XG (a, b), Gal-XG-2 (c, d), and β-6 Galactan-3 epitopes (e, f), are labeled more intensely in roots of space-grown seedlings than in roots of ground controls. Values in box plots in b, d, and f are means of relative fluorescence intensity from randomly picked 10 × 10 μm regions that had positive fluorescence. Box limits indicate 25th and 75th percentiles, horizontal line is the median, and whiskers display minimum and maximum values. **P <0.001 indicates statistical significance as determined by Student’s t-test. Each dot represents individual measurement from ten regions of three longitudinal root tip sections. Bar = 50 μm.

**Fig. 5.**
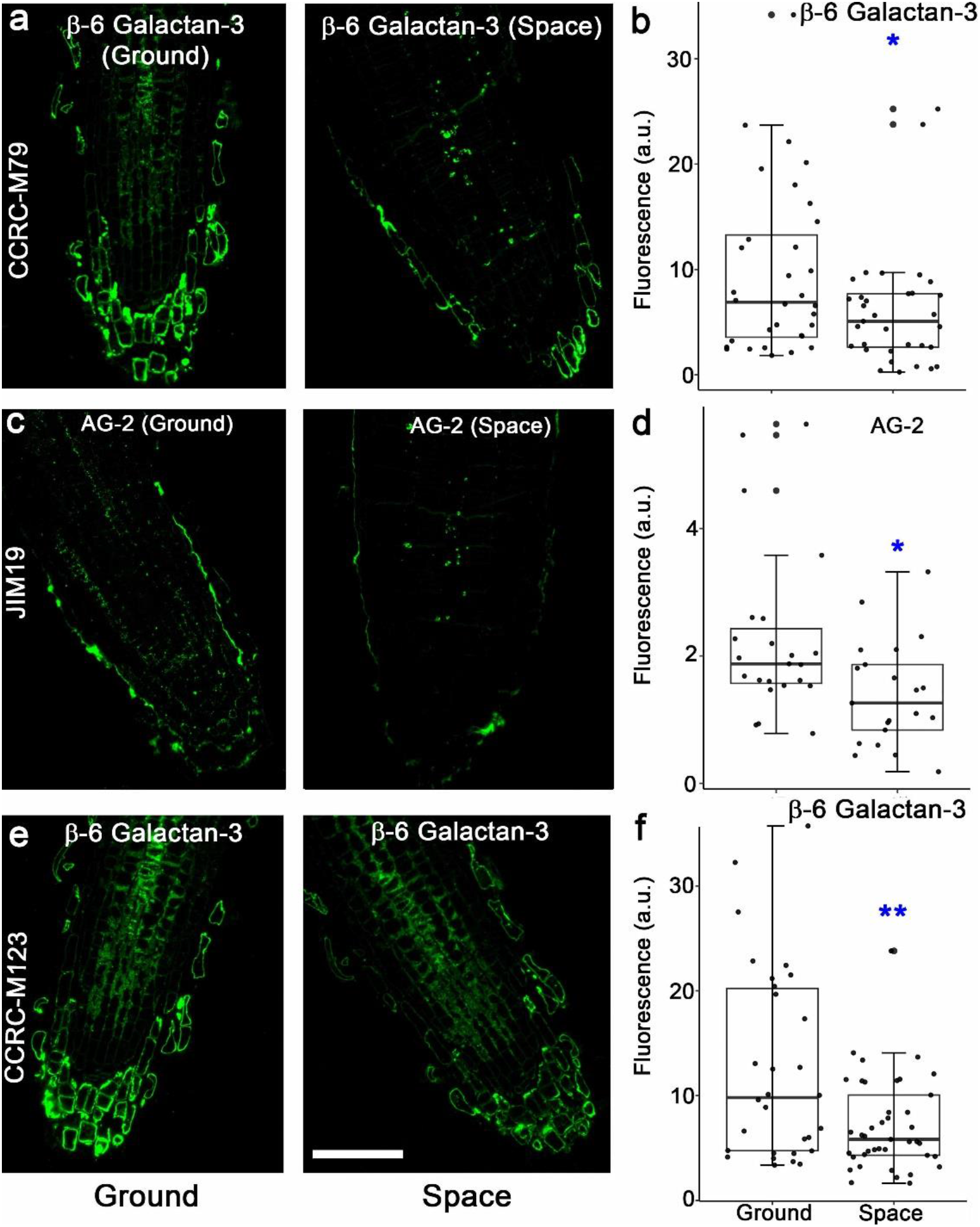
Spaceflight-induced decline in labeling intensity of β-6 Galactan-3 and arabinogalactan (AG)-2 epitopes in root tip longitudinal sections. Epitopes labeled by CCRC-M79 (a, b), JIM19 (c, d), and CCRC-M123 (e, f) had lower fluorescence intensity in roots of 11-day-old seedlings grown in space than that in the ground controls. Values in box plots in b, d, and f are means of relative fluorescence intensity from randomly picked 10 × 10 μm regions that had positive fluorescence. Box limits indicate 25th and 75th percentiles, horizontal line is the median, and whiskers display minimum and maximum values. **P <0.001 and * P<0.01 indicates statistical significance as determined by Student’s t-test. Each dot represents individual measurement from 50 regions of three longitudinal root tip sections. Bar = 50 μm.

Immunohistochemistry of cross-sections of the root maturation zone also revealed spaceflight-induced changes in glycan epitope labeling intensities. Like space-grown root tip longitudinal sections, CCRC-M1 and CCRC-M50, which recognize a fucosylated and galactosylated xyloglucan, respectively, showed space-induced fluorescence increases in root cross sections (Supplementary Fig. 4). Antibodies that recognized Gal-XG-2 (CCRC-M55), M2-XG (CCRC-M99), and 2-Ara-Xylan (CCRC-M108) had lower fluorescence signals in space root cross section than those in ground controls. Four mAbs that bind to xyloglucans (CCRC-M58, CCRC-M96, CCRC-M84, and CCRC-M86) and three β-6 Galactan-3 epitopes (CCRC-M13; CCRC-M79, and CCRC-M123) had fluorescence signals that were similar in root cross section of space- and Earth-grown seedlings (Supplementary Fig. 4 and 5; Fig. 6a).

**Fig. 6.**
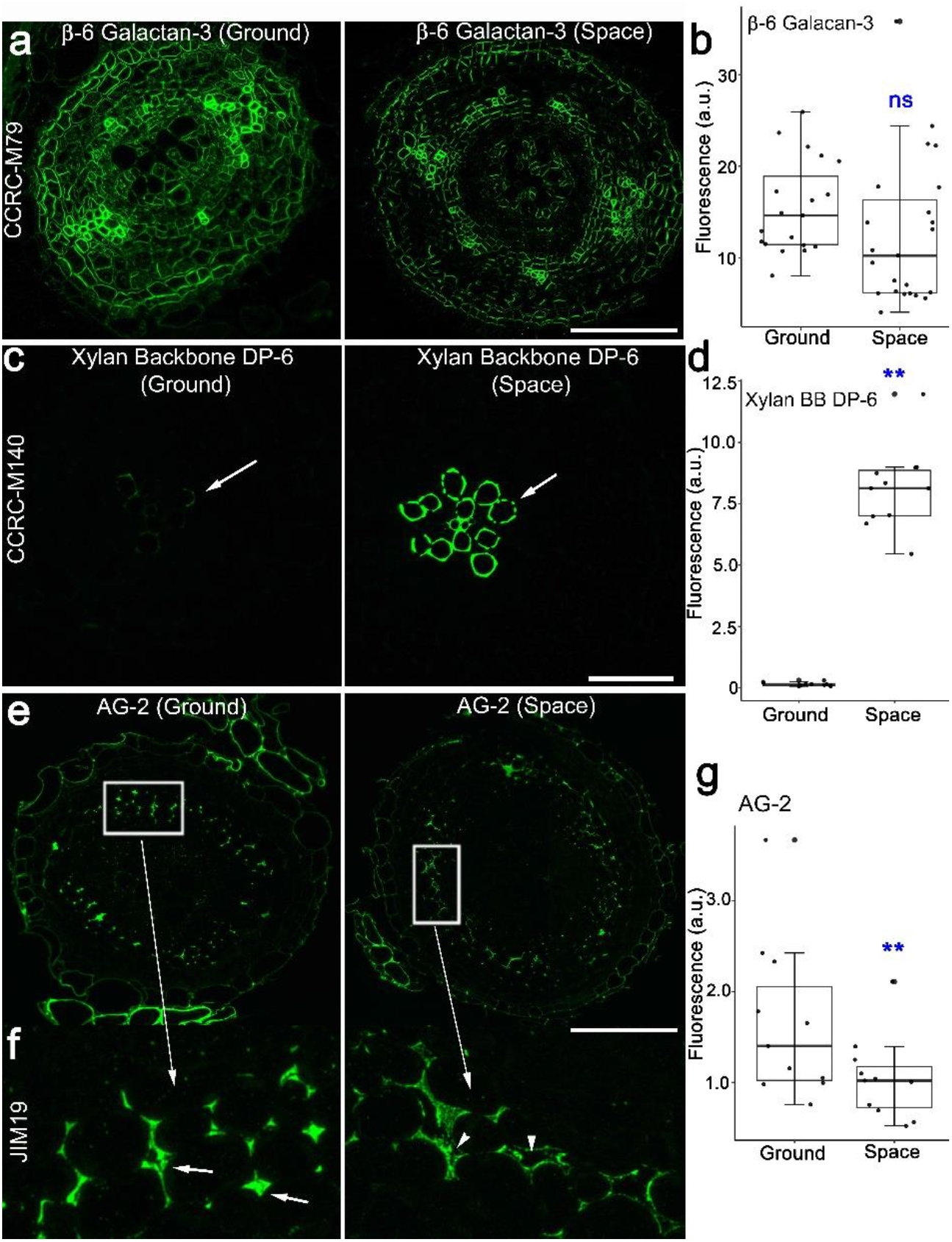
Immunohistochemistry of root cross sections of 11-day old seedlings grown in space and on the ground. (a, b) The β-6 Galactan-3 epitope recognized by CCRC-M79 labels root cell walls uniformly in space and on Earth. (c, d). The xylan backbone mAb, CCRC-M140, preferentially labels root xylem cells in space and on Earth (arrows). Note that xylem cells of roots from space-grown seedlings are more intensely labeled than those of the ground controls (arrows). (e) At low magnification, the AG-2 mAb, JIM19, appears to label root cells of space-grown seedlings in a similar manner as those of seedlings on the ground. (f) Enlarged image of the white boxes in the images in (e). Note that wall spaces in roots of seedlings on the ground have AG-2 epitope fluorescence signals of the that fill the cell wall spaces (arrows). By contrast, cell wall spaces of roots of space-grown seedlings are less densely labeled (arrowheads). () Quantification of fluorescence intensity in root cross sections (b,d, g). Values in box plots are means of relative fluorescence intensity from randomly picked 10 × 10 μm regions that had positive fluorescence. Box limits indicate 25th and 75th percentiles, horizontal line is the median, and whiskers display minimum and maximum values. **P <0.001 indicates statistical significance as determined by Student’s t-test. Not significant (ns). Each dot represents individual measurement from ten regions of three root cross sections. Bar = 50 μm.

Some mAbs that did not label root tip longitudinal sections from space- and Earth-grown seedlings showed positive signals in root cross sections of the maturation zone. Most notable were those that bind to the unsubstituted xylan backbone, such as, CCRC-M138, CCRC-M139, CCRC-M140, and CCRC-M148. Xylan backbone epitopes were more intensely labeled in xylem cells of root cross sections from space-grown seedlings than in the equivalent tissues in ground controls (Fig. 6c, d; Supplementary Fig. 5a, b). CCRC-M148, which also recognizes the xylan backbone, preferentially labeled xylem cells in root cross sections. However, unlike CCRC-M138, CCRC-M139, and CCRC-M140, the intensity of fluorescence signal in xylem cells of CCRC-M148-labeled root cross sections was similar in space and the ground controls (Supplementary Fig. 5a).

Immunohistochemistry of root cross sections also uncovered differences in the density of epitope labeling that were triggered by spaceflight. For instance, labeling of the AG-2-recognizing mAb, JIM19 ^36–38^, was less uniform in the cell wall spaces of roots from space-grown seedlings than that of the ground controls, leading to lower overall fluorescence signals (Fig. 6e-g).

### Some xyloglucan-directed monoclonal antibodies label distinct puncta in root tip longitudinal sections that are reminiscent of post-Golgi organelles

Immunohistochemistry also revealed punctate bodies in root tip longitudinal sections labeled with mAbs against xyloglucans. In particular, CCRC-M1, which binds to the α-fucose-(1, 2)- β- galactose structure of xyloglucan ^39^, labeled distinct bodies in roots of spaceflight- and Earth-grown seedlings (Fig. 7a, b). We cannot discount the possibility that the bodies labeled with anti-xyloglucan mAbs are an artifact of extended aldehyde fixation times. The standard aldehyde fixation time for immunohistochemistry with cell wall mAbs is typically 2 hours ^40^. On the other hand, seedlings in APEX-03-1 used for immunohistochemistry remained in aldehyde fixatives for up to 28 days (Supplementary Fig. 1). We therefore fixed 11-day old *A. thaliana* seedlings for 2 hours and compared CCRC-M1 root labeling patterns of sections from optimally fixed roots with those from APEX-03-1. We found that roots fixed for 2 hours in aldehydes also displayed Fuc-XG-positive bodies, suggesting that immunohistochemistry results from roots fixed in aldehyde for extended periods during APEX-03-1 truly reflected the observed mAb labeling patterns (Fig. 7c).

**Fig. 7.**
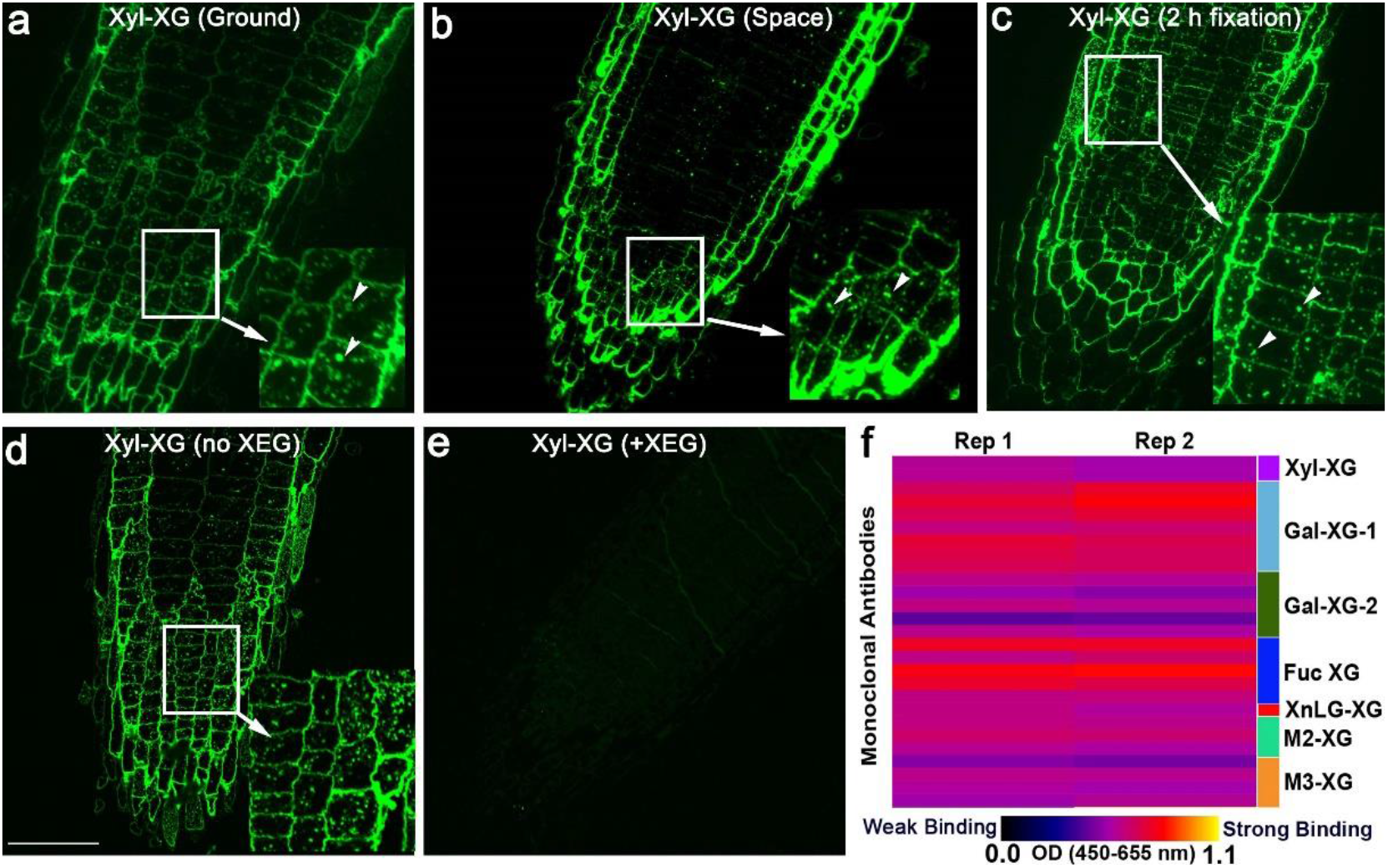
Xyloglucan (XG)-directed mAbs label post-Golgi bodies in root tip longitudinal sections. Root sections from seedlings on the ground (a) and in space (b) labeled with CCRC-M1, which binds to fucosylated xyloglucans (Fuc-XG), have distinct bodies (arrowheads in the enlarged inset). (c) Roots from seedlings optimally fixed for 2 hours in aldehydes also exhibit XG-positive bodies (arrowheads in inset). (d) Without xyloglucan-specific endoglucanase (XEG) pretreatment, CCRC-M1 labels fluorescent bodies in root tip longitudinal sections. (e) Root sections pretreated with XEG, which cleaves off xyloglucan epitopes, followed by CCRC-M1 immunofluorescence labeling, have no fluorescent bodies. (f) Glycome profiling of root microsomal fractions from 11-day-old seedling roots reveal binding of several XG mAbs. Bar in (d) = 10 μm for panels a to e.

The occurrence of fluorescent puncta in root sections labeled with some xyloglucan mAbs was validated further by enzymatic digestion by first treating root sections with a xyloglucan-specific endoglucanase (XEG), which cleaves xyloglucans to oligosaccharides that are released from the tissue sections, prior to immunolabeling. We found that XEG-pretreated roots labeled with CCRC-M1 had no fluorescence (Fig. 7d, e). The bodies labeled with CCRC-M1 are reminiscent of endomembrane-associated Golgi and post-Golgi organelles tagged with fluorescent proteins ^41^. To further verify if xyloglucan mAbs recognize endomembrane components, root microsomal fractions were isolated from 11-day-old *A. thaliana* primary roots and subjected to glycome profiling. Results showed that several xyloglucan mAbs, including CCRC-M1, bound to root microsomal fractions (Fig. 7f).

## DISCUSSION

We demonstrate the use of glycome profiling and immunohistochemistry to better understand how spaceflight affects plant cell wall development in *A. thaliana* seedling roots. We were motivated to implement these methods as part of the APEX-03-1 study because of accumulating transcriptomic studies showing that genes encoding cell wall remodeling proteins are among those that most significantly change expression during spaceflight ^14,16,19,42^. However, information about specific plant cell wall components that are modified by spaceflight is scarce. Unlike Earth-based laboratory experiments, biological spaceflight studies can be fraught with technical challenges because major steps in method implementation are not directly controlled by the investigator team ^43,44^. Furthermore, there are studies indicating that the spaceflight hardware affects plant morphology, as well as gene and protein expression ^45,46^. Adding to these problems is the limited amount of biological material that can be retrieved from spaceflight experiments. Given the technical challenges in conducting plant experiments in space ^47^, it is important to implement procedural controls that can mitigate potential data interpretation errors from biological samples retrieved from space. One procedural control implemented here considered how seedlings were handled during APEX-03-1. This control involved processing *A. thaliana* xyloglucan mutants on the ground in a similar manner as wild-type seedlings grown in space, and asking if glycome profiling still detected cell wall alterations in the mutants despite using only root debris from RNA columns, and subjecting these samples to one step extraction with 4 M KOH. We found that xyloglucan mAb binding to RNA column root debris of the mutants was lower than that of wild type (Supplementary Fig. 2), which was consistent with results obtained from whole plants using the six-step sequential glycome profiling extractions ^34,48^. The procedural control with xyloglucan-defective mutants indicates that glycome profiling of plant debris from RNA purification columns using one extraction step can still reliably report environmentally-induced cell wall changes.

It is also important to note that the aldehydes in APEX-03-1 were stored for an extended period in Kennedy Fixation Tubes (KFTs) prior to seedling fixation, and samples remained in fixative for almost four weeks before they were turned over to the investigator team (Supplementary Fig. 1). Indeed, APEX-03-1 science and payload verification tests (SVT and PVT) revealed that extended aldehyde fixation can adversely affect the quality of root sections ^49^.The results obtained from SVT and PVT enabled us to implement methods for APEX-03-1 that mitigated the extended fixative storage and seedling fixation times. The methods implemented included bubbling the aldehyde solution in nitrogen gas prior to loading in KFTs and storing aldehyde-containing KFTs in the dark and at 4 °C prior to seedling fixation. Furthermore, SVT and PVT indicated that paraformaldehyde and glutaraldehyde in combination could better preserve root morphology than glutaraldehyde alone when seedlings are fixed for long periods ^49^.

Some of the immunohistochemical labeling results presented here were also validated by a post-flight procedural control, in which APEX-03-1-processed roots were compared with roots fixed under standard ground laboratory conditions that involved short duration (*i.e*., 2 hour) seedling fixation and glycome profiling of root membrane fractions (Fig. 7). This control coupled with pretreatment of root sections with xyloglucan digesting enzymes enabled us to show that bodies that were detected by the xyloglucan-directed mAb, CCRC-M1, were likely endomembrane structures. Our results are consistent with membrane fractionation studies demonstrating that CCRC-M1 distributed in the same fractions as proteins that are known to be associated with post-Golgi vesicles ^50^. Moreover, the enrichment of xyloglucan epitopes in the Golgi and *trans*-Golgi Network revealed by glycome profiling of isolated vesicles is consistent with the punctate staining patterns in roots observed here ^51^. It is notable that labeling intensity of xyloglucan-directed mAbs was generally higher in roots from spaceflight seedlings than in roots of the ground controls. It is possible that the spaceflight-induced increases in fluorescence intensity observed after labeling with xyloglucan mAbs is caused by altered biosynthesis and polysaccharide secretion in microgravity. In this regard, proteomic experiments with *A. thaliana* seedlings on the European Modular Cultivation System (EMCS) on-board the ISS uncovered alterations in the expression of membrane-associated proteins by microgravity ^52^. Furthermore, some proteins of *A. thaliana* callus in microgravity were found to be involved in processes related to intracellular trafficking ^53^.

This study highlights the need to validate glycome profiling with immunohistochemistry to gain more meaningful insights into spaceflight-induced modifications in plant cell walls. This is because glycome profiling is done at the whole organ level, while immunohistochemistry focuses on specific locations within the plant organ. Therefore, it is not surprising that results from glycome profiling will not always correlate with immunohistochemistry. Here, validation becomes even more important because only trace amounts of root debris and one extraction step was used for glycome profiling. It was shown previously that such an approach can yield very subtle differences between treatments, and are therefore difficult to interpret without corresponding immunohistochemical studies ^19^. Nonetheless, about 50% of the mAbs selected for immunohistochemistry mirrored results from glycome profiling (*i.e*., fluorescence signals in root sections increased in space), supporting the validity of our approach. The importance of combining glycome profiling and immunohistochemistry on different regions of the plant organ can be seen from results with xylan backbone-directed mAbs. Labeling of xylan backbone epitopes was absent in root tip longitunal sections because xylans are synthesized and deposited in mature tissue, particularly in secondary walls of developing xylem ^54^. This is consistent with the enrichment of fluorescence signals in the xylem in mature root cross sections labeled with xylan-directed CCRC-M138, CCRC-M139, and CCRC-M140 mAbs. By conducting immunohistochemistry in two root regions, one can conclude that spaceflight-induced increases in xylan backbone epitopes uncovered by glycomics are likely the result of localized changes in the xylem. The spaceflight-induced increases in labeling of xylan backbone epitopes in the root xylem with three different mAbs, along with the procedural controls noted earlier, further validates the results presented here. It is unclear why levels of xylan backbone epitopes in the root xylem of space-grown seedlings were higher than those of seedlings on the ground. One possibility is that spaceflight-associated stresses trigger activation of root developmental pathways that are accompanied by hemicellulose and lignin reinforcement of secondary cell walls ^55^. In this regard, it is notable that transcriptome signature of spaceflight-grown *A. thaliana* seedlings overlap with those of cold, drought, and hypoxia ^14^.

The spaceflight-induced root changes in xylans and xyloglucans revealed by glycome profiling and immunohistochemistry are consistent with results showing that the hemicellulose fraction of rice roots increased in space when compared with that of the ground controls ^56^. By contrast, rice shoots on the ISS were shown to have reduced hemicellulose relative to their *1-g* controls ^57^. Differences in microgravity-triggered hemicellulose changes between roots and shoots noted in the above studies further underscore the need for coupling cell wall metabolic analyses with immunolocalizations in different organ- and/or tissue-types.

Our study also reinforces previous work showing a close relationship between spaceflight-induced changes in gene expression and plant phenotypes in space. For instance, shorter root hairs of *A. thaliana* seedlings grown in the Biological Research in Canisters 16 (BRIC-16) hardware onboard the space shuttle Discovery correlates with lower expression of genes encoding cell wall cross-linking class III peroxidases. Some loss-of-function peroxidase mutants grown on Earth have short and ruptured root hairs that are reminiscent of wild-type seedlings grown in space^14^. Consistent with these findings is the reduced formation of ferulate networks, and lower expression of genes encoding class III peroxidases in rice seedlings developing in microgravity^58^. Here, spaceflight-induced changes in xyloglucan epitopes uncovered by glycomics and immunohistochemistry could be the result of differential gene expression. Several transcriptomic studies show that genes encoding xyloglucan endotransglucosylases and glycosyl hydrolases are differentially regulated by spaceflight ^14,16,17,19,59,60^. A RNA-Seq spaceflight study called APEX 03-2 that was conducted at the same time and with the same Veggie growth parameters as our APEX 03-1 is most relevant. Like APEX 03-1, *A. thaliana* roots on APEX 03-2 exhibited a skewing behavior ^42^ (Fig. 1). Results of the APEX 03-2 study showed a number of genes that were differentially regulated in space, including those encoding xyloglucan endotransglucosylases and AG proteins ^61^. RNA-Seq analysis of APEX 03-1 seedling material from which root extracts for the glycomics work described here are ongoing to determine if transcripts differentially induced in space correlate with observed cell wall changes.

In summary, the results presented here support previous studies showing that spaceflight triggers alterations in plant cell walls. These conclusions are based on global differences in binding intensities of mAbs to non-cellulosic glycans from glycome profiling and corresponding immunohistochemical studies of root sections. Given the technical challenges associated with conducting plant spaceflight experiments, this study further highlights the importance of implementing procedural controls that take into account potential data interpretation errors and sample processing artifacts.

## METHODS

### Preflight, flight, and post-flight operations

The experimental timeline of APEX 03-1 is shown in Supplementary Fig. 1, and photographs of various stages involved in the process are shown in Supplementary Fig. 6. Processing of plant material for microgravity experiments was conducted at the Space Station Processing Facility (SSPF) of the NASA Kennedy Space Center (KSC) in Cape Canaveral, Florida in January 2015. Sterilization and planting methods of *A. thaliana* seeds are described in detail in Chai et al. (2022)^62^. Briefly, sterilized seeds of wild-type (ecotype Columbia-0) were planted on growth media (1.2% phyta-agar) consisting of 0.5X Murashige-Skoog (MS) salts and 1% (w/v) sucrose (pH of 5.7) layered onto 90 × 90 mm square Petri dishes (Simport Scientific Inc., Beloeil, Quebec, Canada) (Supplementary Fig. 6a).

Immediately after planting, Petri dishes containing the seeds were exposed to far-red (FR) light (3.2 μmol m^-2^ s^-1^) for 10 minutes using a specially constructed rectangular metal box as described in Nakashima et al. (2014)^49^. Such FR treatment promotes dormancy in *A. thaliana* seeds, preventing premature germination prior to arrival in orbit. A single light emitting diode (LED) (ER-R2, 4606, Norlux Corporation, Carol Stream, Illinois, USA) was used to provide FR illumination to the seeds inside the Petri dishes. FR treatment was carried out under dim light with the distance between the FR LED and Petri dish set at 70 mm. Petri dishes were immediately wrapped with a layer of aluminum foil after FR treatment to keep seeds in complete darkness during imbibition. Petri dishes were then stored in a 4 °C refrigerator prior to launch. The Falcon 9 rocket carrying the Dragon Spacecraft on the SpaceX Commercial Resupply Service (CRS)-5 mission was launched from the Cape Canaveral Air Force Station at 4:47 A.M. (EST) on January 10, 2015 (Supplementary Fig. 6b).

To activate the experiment and trigger germination while in orbit, the aluminum foil was removed, and the Petri dishes were installed in the Veggie plant growth hardware. One Veggie unit was located at the ISS Environmental Simulator (ISSES) located at the SSPF and used as the ground controls (Supplementary Fig. 6c). The other unit was located in the Columbus module on the ISS and used as the spaceflight set (Supplementary Fig. 6d). LEDs on Veggie were programmed to provide constant white light at an intensity of 120-140 μmol m^-2^ s^-1^. Temperature in the Veggie was maintained at 24 °C. After the Petri plates were installed in Veggie, seedlings were harvested at 6 and 11 days after experiment activation and transferred to Kennedy Fixation Tubes (KFTs), containing either RNA*later* (Thermo Fisher Scientific, Waltham, MA) or 4% (v/v) paraformaldehyde and 2.5% (v/v) glutaraldehyde (both from Electron Microscopy Sciences, Hatfield, PA, USA) in phosphate buffered saline (PBS, pH 7.2). Harvesting and fixation on KFTs was performed by Expedition 42 commander Barry Wilmore who completed the process in 2 hours (Supplementary Fig. 6e). KFTs containing the chemicals were kept in the dark and at 4 °C until fixation. Details on KFT actuation are described in Nakashima et al. (2014)^49^. Actuated KFTs with *RNAlater* were stored in a −80°F laboratory freezer (ground controls), or in the Minus Eighty-Degree Laboratory Freezer for ISS (MELFI), whereas those with aldehydes were kept at 4°C. Samples returned to Earth at 4:44 P.M. (PST) on February 11, 2015 (Supplementary Fig. 6f). Fixed seedlings were retrieved from the KFTs at KSC four days after the Dragon Spacecraft splash down and were turned over to the investigator team three days later. Individual seedlings fixed by paraformaldehyde and glutaraldehyde mixture were then carefully separated into shoots and roots, and photographed prior to processing for microscopy.

### Glycome profiling

Seedlings from spaceflight and corresponding ground controls fixed in RNA*later* were thawed, dried using Kimwipes, ground in liquid nitrogen, and stored in −80°F prior to RNA isolation. Seedlings were divided into shoots and roots, and total RNA was isolated using Plant RNeasy Mini Kit (QIAGEN GmbH, Hilden, Germany). The pellet of cell-debris in the QIAGEN shredder spin columns were saved and processed for glycomics.

Glycome profiling of cell wall extracts was conducted using a single step extraction. Briefly, alcohol insoluble materials from root debris were first isolated and extracted with a 4 M KOH solution containing 1% (w/v) sodium borohydride as described in Pattathil et al. (2012)^32^. The extract was dialyzed exhaustively against water and lyophilized before subjecting to glycome profiling. The 4 M KOH extract was then screened by ELISA using a comprehensive suite of plant cell wall glycan-directed mAbs that recognized most of the major non-cellulosic glycan epitopes in plant cell walls^32^. ELISAs were performed by coating 0.3 μg glucose equivalents of carbohydrate materials onto the each well of a 384-well dish (Costar 3700, Corning Inc, Corning, NY, USA). ELISAs were conducted with a fully automated robotic system (Automated ELISA Workstation, Thermo Fisher Scientific Inc. Waltham, MA, USA). The ELISA response values are depicted as heat maps to reflect the relative abundance of glycan epitopes recognized by the mAbs.

### Immunohistochemistry

Immunohistochemistry was performed on roots of 11-day-old seedlings fixed in aldehydes. Roots were washed six times in PBS, dehydrated in a graded ethanol series, and embedded in LR White resin (London Resin Co. Ltd., Reading, Berkshire, UK) as described in Avci et al.(2012)^29^. The resin was then polymerized at 4°C under UV light for 3 days using a PELCO UVC3 Cryo Chamber (Ted Pella, Redding, CA, USA). Serial semi-thin sections (0.25 μm thick) were cut with a diamond knife mounted on a Leica EM UC7 ultramicrotome (Leica Microsystems GmbH, Vienna, Austria). Longitudinal and cross sections were obtained from root tips and the root maturation zone using the flat embedding method described in Avci & Nakashima (2022)^40^. The quality of sections was evaluated by staining in 1% (w/v) Toluidine Blue O in 1% (w/v) sodium borate for 5 min and followed by observation under either a Nikon Microphot-2 microscope (Nikon Corporation, Tokyo, Japan) or a Carl Zeiss ApoTome.2 microscope (Carl Zeiss Microscopy GmbH, Oberkochen, Germany).

Immunohistochemistry was performed by applying and removing a series of 10 μL droplets of the appropriate reagents to the sections as follows: 1). Sections on glass slides were blocked with 3% (w/v) nonfat dry milk diluted in KPBS [0.01 M Potassium Phosphate (pH7.1) containing 0.5 M sodium chloride and 2 mM Sodium Azide (Sigma-Aldrich, St. Louis, Missouri, USA) for 15 minutes; 2). Sections were then incubated with hybridoma supernatant of mAbs to 22 selected cell wall-derived glycans (CarboSource Services, Athens, Georgia, USA) diluted 1:5 in KPBS for 16 to 18 hours at 4 °C; 3). After primary mAb treatment, sections were washed with KPBS three times for 5 minutes each, and incubated for 2 hours at room temperature in the dark in goat anti-mouse IgG conjugated with Alexa-Fluor 488 or goat anti-rat IgG conjugated with Alexa-Fluor 488 (Molecular Probes, Life Technologies Corporation, Carlsbad, California, USA) diluted 1:100 in KPBS; 4). Sections were washed again with KPBS three times for 5 minutes and distilled water for an additional 5 minutes; 5). Finally, sections were mounted on coverslips using Citifluor antifade AF1 (Electron Microscopy Sciences, Hatfield, Pennsylvania, USA). The sections were imaged using a Leica TCS SP2 AOBS Confocal Laser Scanning Microscope (Leica Microsystems CMS GmbH, Mannheim, Germany) equipped with a 40x water immersion objective. Alexa Fluor 488 was imaged by exciting the sections with the 488 nm line of an Argon laser and emission detected at 520 nm using a narrow band pass filter. Magnification and imaging parameters (*i.e*., laser power, pinhole size, and detector sensitivity) were kept constant for flight and ground control samples.

Immunohistochemistry was performed on three sections from the same embedding block that came from three roots for each mAb and treatment. Quantification of fluorescence intensity was conducted by drawing a 10 × 10 μm region of interest in areas of the sections with positive signal using Image J.

### Isolation of root microsomal fractions and xyloglucan-specific endoglucanase treatment

Fresh roots (2 grams) from 11-day-old seedlings were homogenized in 4 mL of a buffer containing 12% (w/w) sucrose, 100 mM Tris/HCl (pH 7.8), 1 mM EDTA, and protease inhibitor cocktails (Sigma-Aldrich, St. Louis, Missouri, USA) as described in Gomez and Chrispeels, (1994)^63^. Cell walls were removed by centrifugation at 1000 × g for 5 minutes. Supernatants were layered over a two-step discontinuous gradient of 16% (w/v) sucrose (5 mL) on top of 48% (w/v) sucrose (1 mL) in 100 mM Tris/HCl (pH7.8) with 1 mM EDTA. The gradient was centrifuged at 150,000 × g for 2 hours and organelles were collected at the 16%/48% interface. Separation of organelles was then performed with an isopycnic 18-50% (w/v) sucrose gradient prepared in 100 mM Tris/HCl (pH7.8) with 1 mM EDTA and centrifuged at 150,000 × g for 2 hours. After centrifugation, 0.6 mL fractions were collected and aliquots were subjected to glycome profiling.

Prior to the immunohistochemistry, root tip longitudinal sections were treated with Xyloglucanase (GH5); E-XEGP (xyloglucan-specific endo-β-(1→4)-glucanase (Megazyme, Wicklow, Ireland) by adding 10 μl endoglucanase solution (0.4U/μl, dissolved in 50 mM Ammonium formate, pH4.5) as described in detail in Günl, et al. (2010)^64^. Sections were incubated in the humid chamber for 16 hours at 37°C followed by washing with Ammonium formate buffer (pH4.5) three times for 5 min. Subsequent immunofluorescence labeling was carried out as described above.

## DATA AVAILABILITY

Data generated in this study are included in this article as a supplemental table and available from the authors on reasonable request.

## ACKNOWLEDEMENTS

This work was supported by NASA grants NNX12AM94G (to E.B.B.), 80NSSC18K1462 and 80NSSC22K0024 (to S.G. and S.C.). Plant cell wall glycan-directed antibodies used here were generated with funding from the National Science Foundation Plant Genome Program (DBI-0421683, to M.G.H.). Glycome profiling of samples described here was supported by a grant to MGH from the NSF Plant Genome Program (IOS-0923992). We thank Allison Mjoen, Howard Levine, Bryan Onate, Trent Smith, Gioia Massa, Colleen Huber, John Carver, Shawn Stephens, Gerald Newsham, David Reed, and Stacy Engel at NASA-KSC for technical support during ground and spaceflight operations.

## AUTHOR CONTRIBUTIONS

J.N., S.P., and U.A. conducted glycome profiling and immunohistochemistry. J.N., S.P., S.C., and E.B.B. analyzed the data, performed statistical analysis, and generated figures. J.N., J.A.S, and E.B.B. prepared seedlings for spaceflight and processed seedlings postflight. S.G., M.G.H., and E.B.B. supervised the project and acquired funding support. All authors contributed to writing and editing the manuscript.

## ADDITIONAL INFORMATION

The online version contains supplementary material.

## Supplementary data legends

**Supplementary Table 1** Raw ELISA readout of monoclonal antibodies (mAbs) used to generate the heatmaps shown in Fig. 2a and Fig. 2b. Each column representing one biological replicate is an average of two technical replicates. The average of three biological replicates in this table were used to plot the bar graphs shown in Fig. 2c.

**Supplementary Fig. 1** Schematic diagram of the experimental timeline of <u>A</u>dvanced <u>P</u>lant <u>EX</u>periments (APEX) 03-1. Note that the *RNAlater* and aldehydes were stored for 19 and 24 days in Kennedy Fixation Tubes (KFTs) prior to seedling fixation on the International Space Station. The experiment was activated by transferring square Petri dishes from 4 °C and darkness to the Veggie unit under continuous white light and 23 °C. Exposure to white light and higher temperatures triggered seeds to germinate on orbit and the ground. Petri dishes were kept in a vertical orientation in Veggie. Fixation was done at 6 and 11 days after experiment activation by transferring seedlings to KFTs with 4% paraformaldehyde and 2.5% glutaraldehyde or RNA*later*. KFTs with aldehydes and RNA*later* were stored at 4 °C and −80 °C, respectively. Seedlings fixed at 6 and 11 days were returned at 28 and 23 days, respectively, for processing.

**Supplementary Fig. 2** Glycome profiling of root cell wall extracts of two xyloglucan mutants from QIAshredder spin column and processed with the single 4M KOH step. Cell wall extracts from the *xxt1/xxt2* double mutant, which makes no xyloglucan, and *mur3-3* single mutant, which lacks galactose-fucose xyloglucan side chains, have lower binding affinity to xyloglucan mAbs (arrowheads) than extracts of wild type seedlings.

**Supplementary Fig. 3** Quantification of root tip longitudinal section fluorescence from space- and Earth-grown seedlings labeled with mAbs to non-cellulosic glycans. Box limits indicate 25th and 75th percentiles, horizontal line is the median, and whiskers display minimum and maximum values. ***P<0.0001, **P <0.001, and *P<0.01 indicate statistical significance as determined by Student’s t-test. Not significant (ns). Each dot represents individual measurement from 30-50 regions of three root tip longitudinal sections. Xyloglucan (XG); Galactose (Gal); Fucose (Fuc); Arabinose (Ara); Xylose (Xyl).

**Supplementary Fig. 4** Quantification of root cross section fluorescence from space- and Earth-grown seedlings labeled with mAbs to non-cellulosic glycans. Box limits indicate 25th and 75th percentiles, horizontal line is the median, and whiskers display minimum and maximum values. ***P<0.0001, **P <0.001, and *P<0.01 indicate statistical significance as determined by Student’s t test. Not significant (ns). Each dot represents individual measurement from 20-30 regions of three root cross sections. Xyloglucan (XG); Galactose (Gal); Fucose (Fuc); Arabinose (Ara); Xylose (Xyl).

**Supplementary Fig. 5** Immunohistochemistry of root cross sections from space- and Earth-grown seedlings labeled with mAbs to non-cellulosic glycans. (a) Fluorescence of root cross sections labeled with CCRC-M123 and CCRC-M148 were not significantly different statistically between space and ground controls. Root cross sections from seedlings grown in space labeled with CCRC-M138 and CCRC-M139 had higher fluorescence than those of the ground controls. Box limits indicate 25th and 75th percentiles, horizontal line is the median, and whiskers display minimum and maximum values. ***P<0.0001 indicate statistical significance as determined by Student’s t test. Not significant (ns). Each dot represents individual measurement from 20-30 regions of three root cross sections. (b) CCRC-M123 labels roots cells uniformly in space and on Earth. The xylan mAbs, CCRC-M138 and 139, preferentially labels root xylem cells in space and on Earth (arrows). Note that xylem cells of roots from space-grown seedlings labeled with CCRC-M138 and 139 are more intensely labeled than that of the ground controls. Size of the bars are indicated in the figure. Degree of Polymerization (DP).

**Supplementary Fig. 6** Overview of preflight, flight and postflight operations for glycome profiling and immunohistochemistry of *A. thaliana* seedling roots. (a) Planting of *A. thaliana* seeds in square Petri dishes under a laminar flow hood at the Space Station Processing Facility (SSPF) eight days prior to Space X-5 launch. (b) Launch of the SpaceX Falcon rocket carrying the Dragon spacecraft with the Petri dishes containing *A. thaliana* seeds. (c) Veggie hardware at the SPFF used for the ground controls. (d) Veggie hardware located at the Columbus module of the International Space Station (ISS). (e) Harvesting 11-day-old seedlings and transferring them to Kennedy Fixation Tubes (KFTs) containing chemical fixatives. (f) The Dragon spacecraft with the APEX 03-1 *A. thaliana* seedlings being lifted onto the deck of a recovery ship in the Pacific Ocean. Images in panels b, d, e, and f are courtesy of NASA and are in the public domain.

