## Supplementary Figures for "Glycome profiling and immunohistochemistry uncover spaceflight-induced changes in non-cellulosic cell wall components in *Arabidopsis thaliana* seedling roots"

**Supplementary Figures (Nakashima et al)**

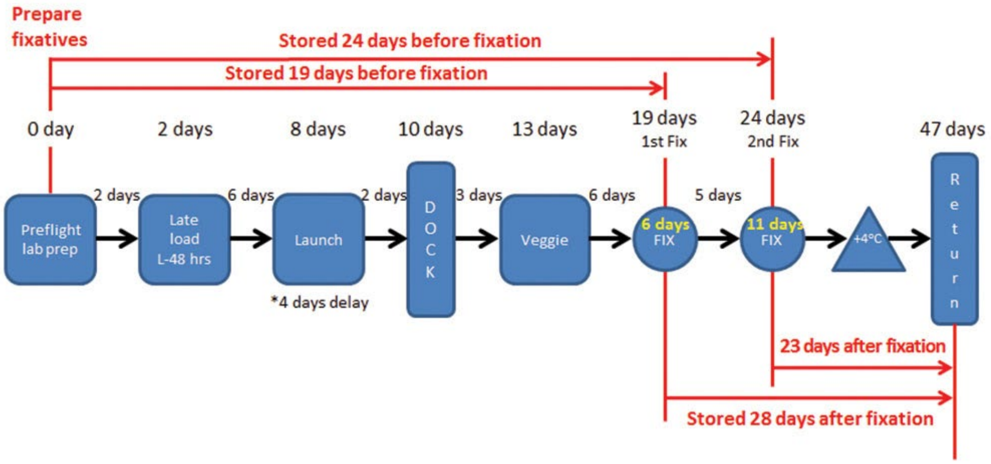


**Supplementary Fig. 1** Schematic diagram of the experimental timeline of Advanced Plant EXperiments (APEX) 03-1. Note that the RNA*later* and aldehydes were stored for 19 and 24 days in Kennedy Fixation Tubes (KFTs) prior to seedling fixation on the International Space Station. The experiment was activated by transferring square Petri dishes from 4 °C and darkness to the Veggie unit under continuous white light and 23 °C. Exposure to white light and higher temperatures triggered seeds to germinate on orbit and the ground. Petri dishes were kept in a vertical orientation in Veggie. Fixation was done at 6 and 11 days after experiment activation by transferring seedlings to KFTs with 4% paraformaldehyde and 2.5% glutaraldehyde or RNA*later*. KFTs with aldehydes and RNA*later* were stored at 4 °C and -80 °C, respectively. Seedlings fixed at 6 and 11 days were returned at 28 and 23 days, respectively, for processing.

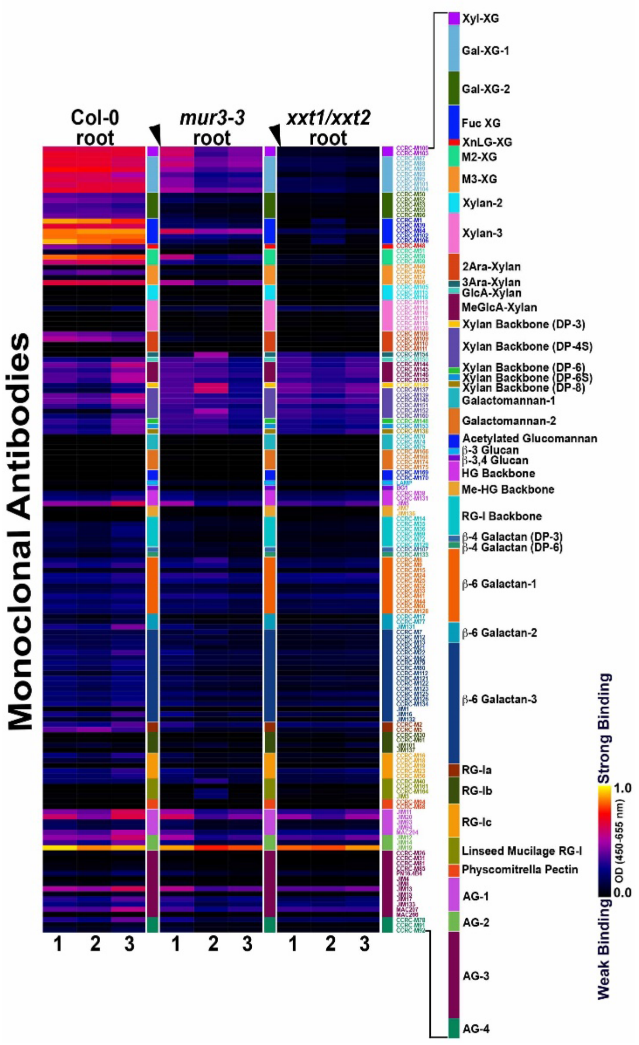


**Supplementary Fig. 2** Glycome profiling of root cell wall extracts of two xyloglucan mutants from QIAshredder spin column and processed with the single 4M KOH step. Cell wall extracts from the *xxt1/xxt2* double mutant, which makes no xyloglucan, and *mur3-3* single mutant, which lacks galactose-fucose xyloglucan side chains, have lower binding affinity to xyloglucan mAbs (arrowheads) than extracts of wild type seedlings.

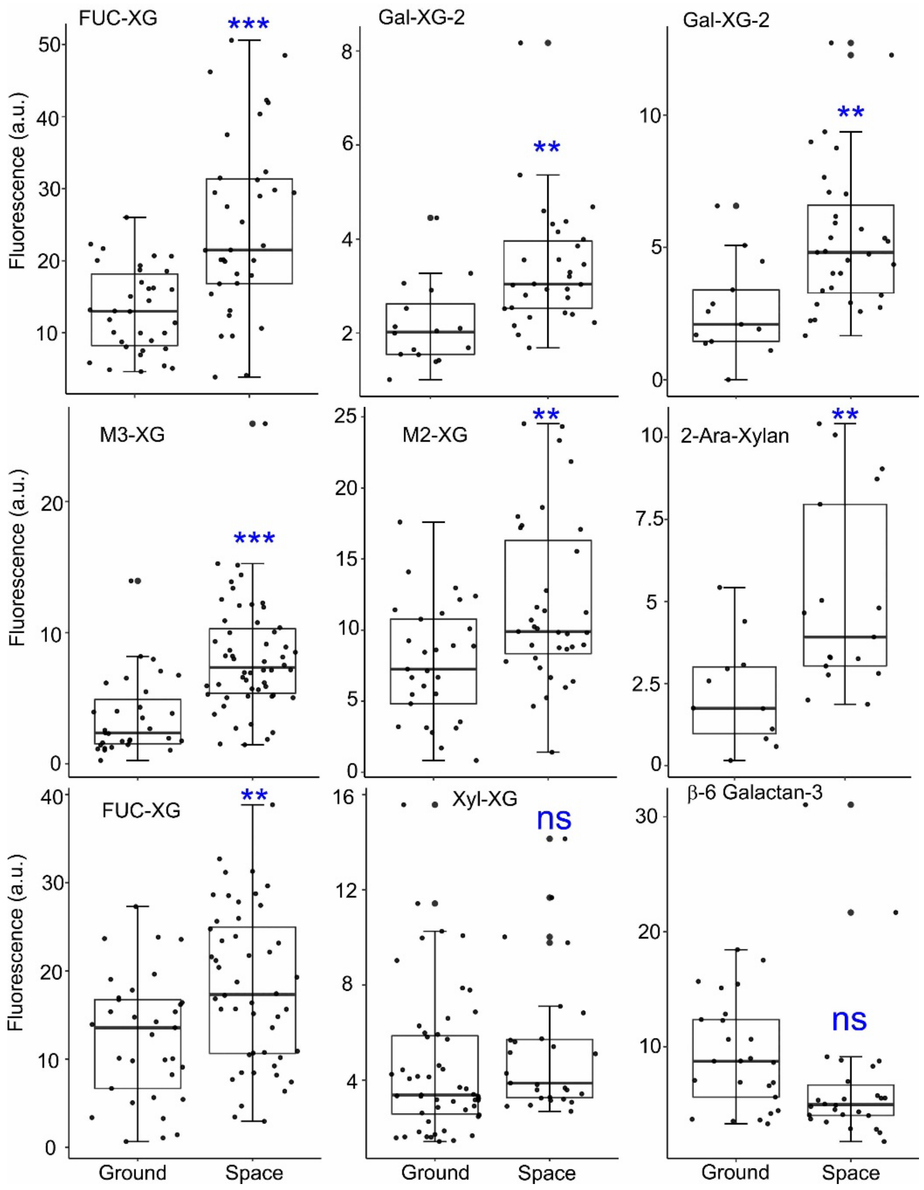


**Supplementary Fig. 3** Quantification of root tip longitudinal section fluorescence from space- and Earth-grown seedlings labeled with mAbs to non-cellulosic glycans. Box limits indicate 25th and 75th percentiles, horizontal line is the median, and whiskers display minimum and maximum values. ***$P<0.0001$, **$P<0.001$, and *$P<0.01$ indicate statistical significance as determined by Student's t-test. Not significant (ns). Each dot represents individual measurement from 30-50 regions of three root tip longitudinal sections. Xyloglucan (XG); Galactose (Gal); Fucose (Fuc); Arabinose (Ara); Xylose (Xyl).

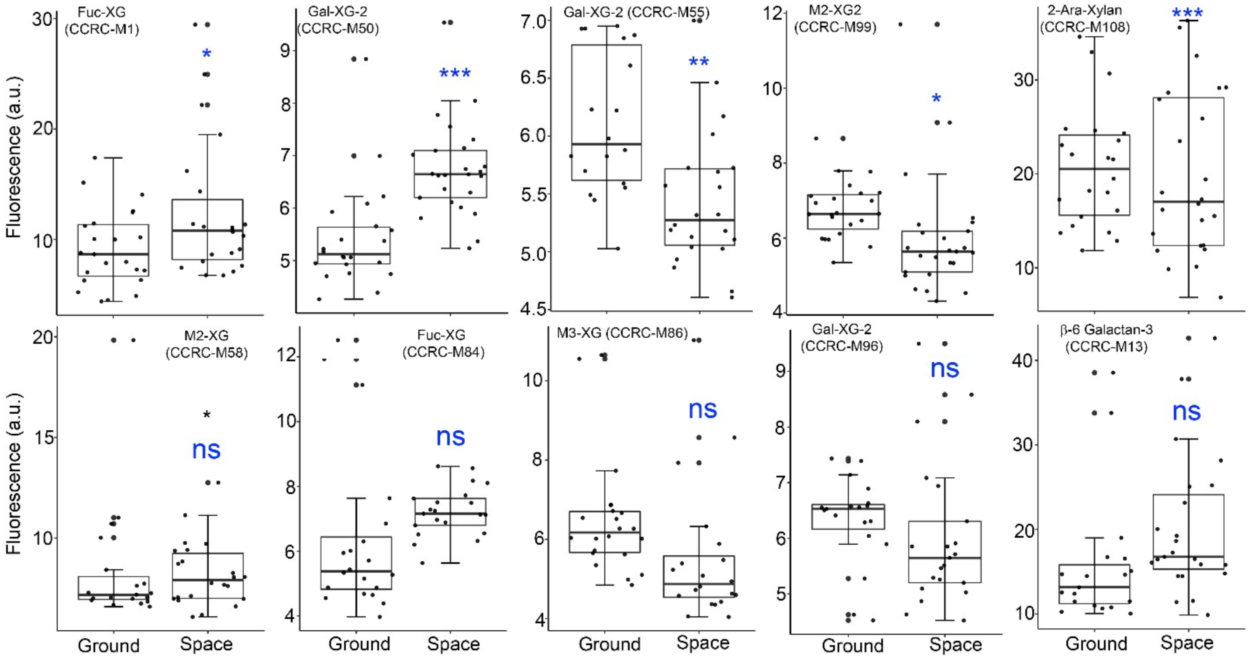


**Supplementary Fig. 4** Quantification of root cross section fluorescence from space- and Earth-grown seedlings labeled with mAbs to non-cellulosic glycans. Box limits indicate 25th and 75th percentiles, horizontal line is the median, and whiskers display minimum and maximum values. ***P<0.0001, **P <0.001, and *P<0.01 indicate statistical significance as determined by Student's t test. Not significant (ns). Each dot represents individual measurement from 20-30 regions of three root cross sections. Xyloglucan (XG); Galactose (Gal); Fucose (Fuc); Arabinose (Ara); Xylose (Xyl).

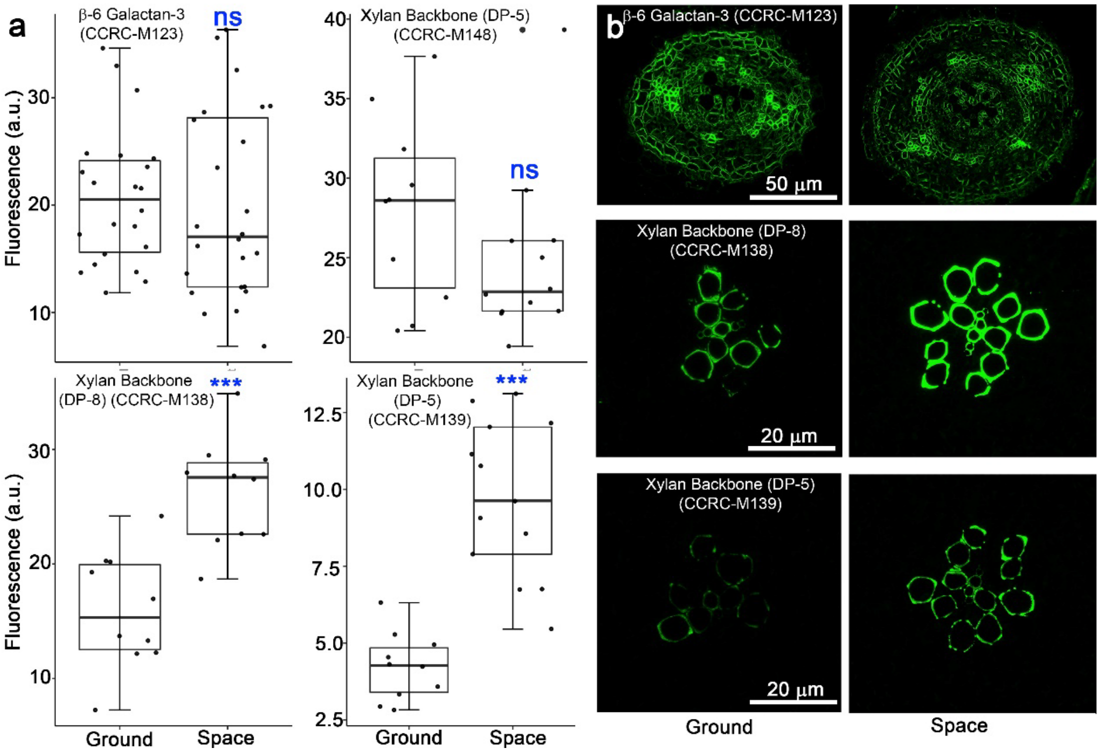


**Supplementary Fig. 5** Immunohistochemistry of root cross sections from space- and Earth-grown seedlings labeled with mAbs to non-cellulosic glycans. (a) Fluorescence of root cross sections labeled with CCRC-M123 and CCRC-M148 were not significantly different statistically between space and ground controls. Root cross sections from seedlings grown in space labeled with CCRC-M138 and CCRC-M139 had higher fluorescence than those of the ground controls. Box limits indicate 25th and 75th percentiles, horizontal line is the median, and whiskers display minimum and maximum values. ***$P<0.0001$ indicate statistical significance as determined by Student's t test. Not significant (ns). Each dot represents individual measurement from 20-30 regions of three root cross sections. (b) CCRC-M123 labels roots cells uniformly in space and on Earth. The xylan mAbs, CCRC-M138 and 139, preferentially labels root xylem cells in space and on Earth (arrows). Note that xylem cells of roots from space-grown seedlings labeled with CCRC-M138 and 139 are more intensely labeled than that of the ground controls. Size of the bars are indicated in the figure. Degree of Polymerization (DP).

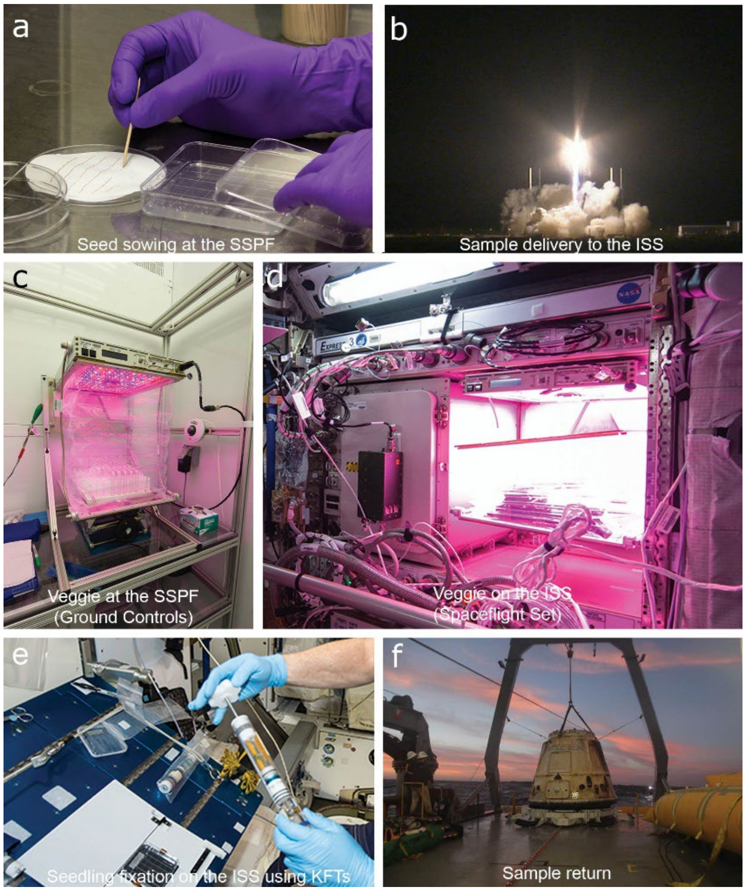


**Supplementary Fig. 6** Overview of preflight, flight and postflight operations for glycome profiling and immunohistochemistry of *A. thaliana* seedling roots. (a) Planting of *A. thaliana* seeds in square Petri dishes under a laminar flow hood at the Space Station Processing Facility (SSPF) eight days prior to Space X-5 launch. (b) Launch of the SpaceX Falcon rocket carrying the Dragon spacecraft with the Petri dishes containing *A. thaliana* seeds. (c) Veggie hardware at the SPFF used for the ground controls. (d) Veggie hardware located at the Columbus module of the International Space Station (ISS). (e) Harvesting 11-day-old seedlings and transferring them to Kennedy Fixation Tubes (KFTs) containing chemical fixatives. (f) The Dragon spacecraft with the APEX 03-1 *A. thaliana* seedlings being lifted onto the deck of a recovery ship in the Pacific Ocean. Images in panels b, d, e, and f are courtesy of NASA and are in the public domain.
